# Universal agentic probe design for imaging-based spatial-omics

**DOI:** 10.64898/2026.04.12.717982

**Authors:** Qian Zhang, Huaiyuan Cai, Jiahao Zhang, Lingkai Zhang, Xiaoyu Wu, Yuting Wei, Yu Chen, Xiaofeng Wu, Wen Su, Wenbao Qi, Xiaojie Qiu, Gang Cao, Weize Xu

## Abstract

Probe design for fluorescence in situ hybridization (FISH) underpins spatial transcriptomics, three-dimensional genome studies, and clinical diagnostics, yet remains constrained by two challenges: dependence on expert knowledge for parameter selection and quality evaluation, and the inability of existing tools to accommodate the diverse probe architectures introduced by rapidly emerging methods. Here we present U-Probe, a universal and agentic probe design platform. U-Probe employs a declarative configuration system with a directed acyclic graph (DAG)-based assembly engine that supports arbitrary probe structures from established protocols such as MERFISH, seqFISH, and MiP-seq to entirely novel architectures without code modifications. Integrated LLM-based AI agents enable conversational design workflows, allowing users to specify experimental goals in natural language and receive synthesis-ready probe sequences. We validated U-Probe in three scenarios: agent-driven MiP-seq panel design for influenza-infected mouse lung tissue, genome-tiling DNA-FISH for herpesvirus detection, and a novel RCA-based ligation probe for single-nucleotide mutation discrimination. U-Probe is available as an open-source tool with CLI, Web, and agent interfaces.

## Introduction

Fluorescence in situ hybridization (FISH)-based probe design underpins a broad and growing range of applications, from spatial transcriptomics^1^ and three-dimensional genome architecture studies^2^ to clinical diagnostics^3^. The demand is particularly acute for non-model organisms, where commercial antibodies are largely unavailable and FISH probes—requiring only genomic or transcriptomic sequences—offer the primary route for in situ gene detection. Single-cell sequencing has further amplified this need by generating transcriptomic atlases across diverse species that now require spatial validation, driving an explosion in the demand for custom probe sets. Yet this field faces two fundamental challenges. First, despite the availability of probe design tools such as OligoMiner^4^, PaintSHOP^5^, Tigerfish^6^, eFISHent^7^, and gene2probe^8^, the design process remains heavily dependent on expert knowledge—users must understand how probe structures match experimental protocols, select appropriate thermodynamic filtering parameters, and evaluate off-target specificity—a labor-intensive process that poses a substantial barrier for groups without dedicated computational expertise. Second, the rapid emergence of novel methods—including PRISM^9^, TDDN-FISH^10^, RAEFISH^11^, OpenFISH^12^, and Oligo-LiveFISH^13^—has diversified probe architectures far beyond what existing tools can accommodate. These tools hard-code support for established protocols like MERFISH^14^ or DNA chromosome painting^15^, forcing laboratories developing novel methods to build custom pipelines from scratch.

Here we present U-Probe, a universal and agentic probe design platform that addresses both limitations. U-Probe enables design of arbitrary probe structures through a declarative configuration system, resolving the architectural fragmentation problem; and integrates large language model (LLM)-based agents for conversational design workflows, lowering the expertise barrier by guiding users through the entire design process via natural language interaction.

## Methods

The U-Probe platform integrates a declarative YAML-based configuration system with a directed acyclic graph (DAG) assembly engine to design arbitrary probe architectures. Quality metrics, including melting temperature, secondary structure stability, and off-target specificity, are computed using established thermodynamic models and alignment tools. The LLM-based multi-agent system is built upon the PantheonOS framework, enabling natural language-driven probe design. Experimental validations, including MiP-seq, DNA-FISH, and padlock probe-based FISH for single-nucleotide mutation detection, were performed using standard protocols. Detailed computational procedures, algorithm descriptions, and experimental protocols are provided in the Supplementary Methods.

## Results

### Declarative configuration system for universal probe design

U-Probe’s universality stems from a general declarative configuration system combined with a directed acyclic graph (DAG)-based assembly engine (Fig. 1a). Users define probes as modular templates with named parts. Each part can reference target sequences (with slicing and reverse complement operations), barcodes from encoding tables, or other probe components. The DAG engine resolves dependencies between parts via topological sort, assembling arbitrarily complex multi-part probes from these declarations. For example, an RCA probe configuration specifies a circle probe with target-complementary arms flanking barcodes, plus an amplification probe referencing both the circle probe and target region. This approach supports DNA-FISH (single locus, chromosome painting^15^, telomere), RNA-FISH (smFISH^16^, MERFISH^14^, seqFISH^17^, *π*-FISH^18^, MiP-seq^19^), and entirely novel architectures without code modifications (Fig. 1b).

**Fig. 1:**
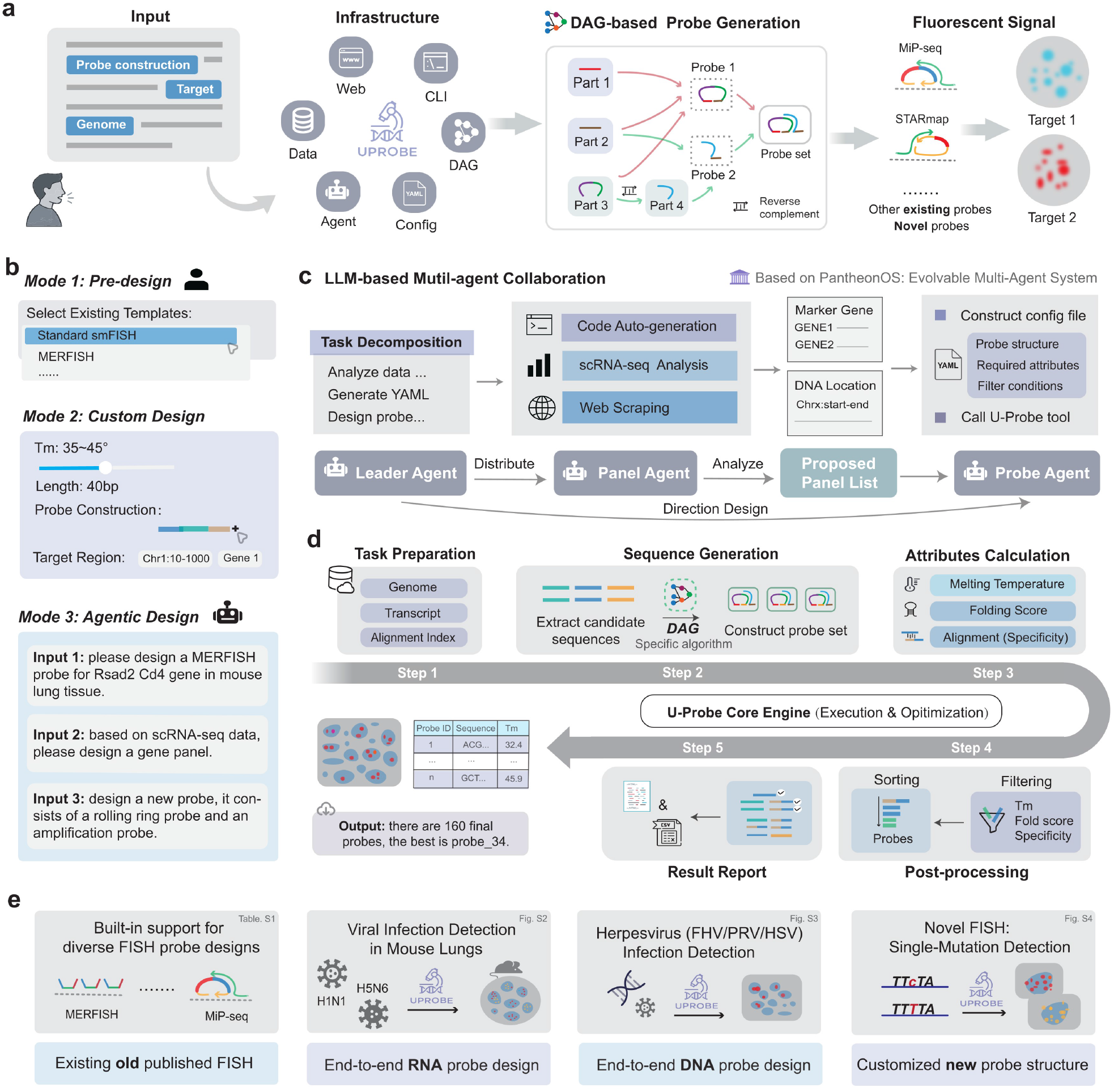
U-Probe: Universal agentic probe design platform. **a**, System architecture. Users interact via Web, CLI, or AI Agent. A DAG-based engine assembles modular probe parts into complete probe sets. **b**, Three design modes: Pre-design (built-in templates), Custom Design (user-defined parameters), and Agentic Design (natural language instructions). **c**, LLM-based multi-agent framework. A Leader Agent coordinates the Panel Agent (gene panel optimization) and Probe Agent (YAML configuration and probe generation). **d**, Core engine pipeline: task preparation, sequence generation, attributes calculation, post-processing, and result reporting. **e**, Application scenarios: built-in FISH protocol support (Supplementary Table. S1), end-to-end RNA probe design (Supplementary Fig. S1), DNA probe design (Supplementary Fig. S2), and novel probe structure design (Supplementary Fig. S3).

Beyond probe structure, the configuration also specifies quality metrics and filtering criteria. U-Probe computes GC content, melting temperature, secondary structure stability (via ViennaRNA^20^), off-target mapping (via Bowtie2^21^), and k-mer frequency (via Jellyfish^22^) for each candidate probe. Users define filtering conditions, sorting priorities, and post-processing steps including overlap removal and equal spacing for tiling designs. Together, this declarative system separates experimental intent from computational execution, enabling researchers to focus on probe design rather than pipeline engineering.

### LLM-based multi-agent system for conversational design

Beyond configuration-based design, U-Probe integrates LLM-based AI agents through the PantheonOS framework^23^ (Fig. 1c). A hierarchical team of agents—Leader, Panel, and Probe—interprets natural language requests and executes multi-step design pipelines. Users can describe experimental goals or novel probe architectures in plain language, and the agents autonomously construct configuration files, select appropriate parameters, and invoke the U-Probe engine (Fig. 1d), enabling rapid prototyping without requiring expertise in probe thermodynamics or configuration syntax. This also bridges single-cell and spatial analyses: researchers can directly provide scRNA-seq data, and the agent will identify marker genes and design the corresponding probe panel for spatial validation.

### Experimental validation of U-Probe

We validated U-Probe across three experimental scenarios. In two agent-driven workflows, the system autonomously designed a MiP-seq probe panel from mouse lung scRNA-seq data for H5N6 and H1N1 influenza-infected tissue sections, successfully resolving spatial cell type distributions and differential immune infiltration patterns (Supplementary Fig. S1); and designed genome-tiling DNA-FISH probe pools for three herpesviruses (FHV, PRV, and HSV), achieving high genome coverage (77–98%) and enabling rapid detection of viral infection in cultured cells (Supplementary Fig. S2). To demonstrate the flexibility of the configuration system, we also designed a novel RCA-based ligation probe for single-nucleotide mutation detection of the H9N2 avian influenza PB2 627E/K polymorphism— a probe architecture that existing tools cannot express—achieving allele-specific discrimination and multiplexed co-infection detection in a single experiment (Supplementary Fig. S3).

### U-Probe Web platform

We implemented U-Probe Web (Supplementary Fig. S4), providing visual exploration of probe structures, form-based batch submission, a conversational agent interface, and synthesis-ready outputs.

## Discussion

By enabling arbitrary probe architectures through declarative configuration and expert-free design through AI agents, U-Probe addresses both the fragmentation of probe design tools and the expertise barrier in imaging-based spatial-omics. Looking forward, U-Probe can automatically interface with genomes and transcriptomes of any species—including polyploid organisms—and the integration with the evolvable PantheonOS agent framework^23^ allows the system to continuously incorporate new design strategies as spatial-omics technologies advance.

## Acknowledgements

We thank Cong Wang for his valuable contributions to this work during his master’s studies. The entire manuscript has been checked for correctness, and the authors are responsible for its final content.

## Author contributions

W.X. conceived the study. W.X., Q.Z., and H.C. led the U-Probe project. Q.Z., W.X., H.C., J.Z., and Y.C. developed the U-Probe core engine. Q.Z. designed and implemented the U-Probe web frontend and U-Probe agent team. Q.Z., W.X., and H.C. designed the main figures. H.C. coordinated the experiments, performed data analyses, and generated the plots. J.Z., L.Z., X.W., and Y.W. performed the experiments. G.C., X.Q., W.Q., X.W., Y.C., and W.S. provided critical advice. W.X. and G.C. supervised the project and secured funding. W.X. wrote the first draft. All authors edited and approved the manuscript.

## Data availability

Example configurations and datasets are available at https://github.com/UFISH-Team/U-Probe.

## Code availability

U-Probe is open-source at https://github.com/UFISH-Team/U-Probe. U-Probe Web frontend: https://github.com/UFISH-Team/uprobe-web-ui.

## Supplementary Information

### Supplementary Methods

#### Declarative probe design via YAML configuration

U-Probe employs a YAML-based configuration system that separates probe specification from implementation. A protocol file defines four key sections: (1) *target extraction*, specifying the genomic source (genome, exon, CDS, or UTR), target sequence length, and tiling overlap; (2) *probe structure*, defining probes as composable templates with named parts that can reference target sequences, barcodes from encoding tables, or other probe components via expression syntax (supporting Python slicing, reverse complement, and variable substitution); (3) *quality attributes*, listing which metrics to compute for each probe part; and (4) *post-processing*, specifying filtering conditions, sorting priorities, overlap removal, and equal-spacing parameters. If quality attributes or post-processing sections are omitted, U-Probe automatically generates sensible defaults based on the probe type (DNA or RNA mode).

#### DAG-based probe generation

Probe assembly follows a directed acyclic graph (DAG) execution model. Formally, a probe design is represented as a DAG *G* = (*V, E*), where each node *v* ∈ *V* corresponds to a probe component (part) and each directed edge (*u, v*) ∈ *E* indicates that node *v* depends on the output of node *u*. Nodes are classified into two types:

- *Expression nodes v*_*e*_: evaluate a formula *f* (*C*) over a context *C* = {*s, t*, **, rc}, where *s* is the target sequence, *t* is the target identifier, ** is the barcode encoding table, and rc(·) is the reverse complement operator. For example, *f* (*C*) = rc(*s*[*i* : *j*]) ∥ **[*t*][*k*] extracts a subsequence, takes its reverse complement, and concatenates a barcode.
- *Template nodes v*_*t*_: concatenate child outputs via a template string *T* = “{*p*_1_}{*p*_2_} …{*p*_*n*_}”, where each *p*_*i*_ refers to a child node. Template nodes may themselves contain child template nodes, enabling recursive composition of arbitrarily complex probe architectures.

The DAG is resolved by topological sort. Let deg^*−*^(*v*) denote the in-degree of node *v*. All nodes with deg^*−*^(*v*) = 0 (leaf nodes) are enqueued first. Each node *v* is evaluated only when all its predecessors {*u* : (*u, v*) ∈ *E*} have completed, after which *v*’s dependents are checked and enqueued if ready. This guarantees correct evaluation order regardless of DAG depth or branching structure.

For a set of *N* target sequences {*s*_1_, …, *s*_*N*_ }, the DAG is instantiated independently for each *s*_*i*_, producing *N* probe records. Each record contains the final assembled probe and all intermediate parts as separate fields, enabling per-part quality assessment in subsequent steps.

#### Quality metrics computation

U-Probe computes the following attributes for candidate probes: *GC content* as the fraction of G and C nucleotides; *melting temperature* (*T*_*m*_) via nearest-neighbor thermodynamics (Primer3); *folding score* as the minimum free energy of secondary structure predicted by ViennaRNA^1^; *self-match score* by counting complementary k-mer pairs between the probe and its reverse complement; *mapped sites* by aligning probes to the reference genome using Bowtie2^2^ in sensitive local mode (reporting up to 100 alignments, MAPQ ≥ 30); *k-mer frequency* by querying a pre-built Jellyfish^3^ index to obtain the maximum genomic occurrence count across all k-mers in a probe; and *number of mapped genes* by aligning probes to a transcript database and counting unique gene hits.

#### Post-processing pipeline

After attribute computation, probes undergo a multi-step post-processing pipeline. First, user-defined *filters* remove probes failing quality thresholds (e.g., GC content outside 30–70%, *T*_*m*_ outside 50–70°C). Second, an optional *off-target peak avoidance* step uses a greedy algorithm to exclude probes with high off-target alignment density within a user-defined genomic window. Third, *overlap removal* greedily selects non-overlapping probes sorted by genomic position, enforcing a minimum inter-probe distance. Fourth, *equal spacing* downsamples probes to a desired count by selecting uniformly distributed indices across the target region. Finally, probes are *sorted* by multiple quality metrics.

#### LLM-based multi-agent system

U-Probe integrates large language model (LLM)-based agents through the PantheonOS multi-agent framework^4^. The system employs a hierarchical team of three specialized agents. The *Leader Agent* parses user requests, routes tasks, and coordinates the overall workflow. The *Panel Agent* performs computational analyses including single-cell RNA-seq processing (clustering, differential expression), web scraping for literature-curated marker genes, and code generation for panel optimization. The *Probe Agent* translates design specifications into YAML protocol files by referencing built-in templates, then invokes the U-Probe core engine to execute probe generation. Agents communicate through structured handoff protocols, enabling end-to-end workflows from raw data to synthesis-ready probe sequences via natural language conversation.

#### U-Probe Web platform

The U-Probe Web platform consists of a REST API backend and a React-based frontend. The platform provides five integrated modules: a *Genome Data Center* for uploading and managing reference genomes (FASTA) and annotation files (GFF/GTF); a *Design Dashboard* offering both guided step-by-step workflows and advanced custom probe type interfaces; a *Task Monitor* for tracking computational progress with real-time status updates; a *System Portal* for platform navigation; and a *Conversational Agent* interface that connects to the LLM-based multi-agent system for interactive probe design with conversation history management. The platform supports user authentication, file management with drag-and-drop upload, and downloadable result reports in HTML and CSV formats.

### Ethical statement, virus isolation and propagation, and animal experiments

All animal experiments were conducted in Animal Biosafety Level 3 (ABSL-3) facilities in strict accordance with protocols approved by the Institutional Biosafety Committee of South China Agricultural University (SCAU). All procedures involving animals were reviewed and approved by the SCAU Institutional Animal Care and Use Committee (IACUC; Approval No. 2017A002).

The H1N1 influenza virus (A/PR/8/34 [H1N1], hereafter PR8/H1N1) and H5N6 influenza virus (A/duck/Guangdong/673/2014 [H5N6], hereafter 673/H5N6) were obtained from the National Avian Influenza Para-Reference Laboratory at SCAU. Viruses were propagated in 10-day-old specific-pathogen-free (SPF) embryonated chicken eggs at 37 °C and stored at −80 °C until use.

Six-week-old female BALB/c mice (Vital River, China) were used for infection experiments. Mice were randomly assigned into experimental groups (*n* = 3 per group) and intranasally inoculated with 10^6^ 50% egg infectious dose (EID_50_) of the respective influenza virus in 50 *µ*L phosphate-buffered saline (PBS). Control mice (*n* = 3) received PBS only. At 3 days post-infection (dpi), mice were euthanized, and lung tissues were aseptically collected. Tissues were embedded in Optimal Cutting Temperature (OCT) compound for subsequent spatial transcriptomic analysis.

### MiP-seq in situ RNA detection

Padlock probes were first phosphorylated at the 5^*′*^ end using T4 Polynucleotide Kinase (Vazyme, N102-01) to enable subsequent MiP-seq reactions. Phosphorylated padlock probes and an equimolar concentration of rolling circle amplification (RCA) initiators (30 nM each) were prepared in hybridization buffer containing 2× SSC (Sangon, B548109), 10% formamide (Sangon, A100606), and 20 mM ribonucleoside–vanadyl complexes (Beyotime, R0108). The mixture was applied to samples (cultured cells or pretreated tissue sections) and incubated overnight at 37 °C.

Following hybridization, ligation was performed using a mixture containing 1 U *µ*L^*−*1^ SplintR ligase (NEB, M0375L), 1× ligation buffer, and 0.2 U *µ*L^*−*1^ RiboLock RNase inhibitor at 37 °C for 2 h. Samples were then washed twice with DEPC-treated PBSTR for 20 min each.

Rolling circle amplification (RCA) was carried out using a reaction mixture containing 1 U *µ*L^*−*1^ Phi29 DNA polymerase (Vazyme, N106-01), 1× RCA buffer, 0.25 *µ*M dNTPs, 0.2 *µ*g *µ*L^*−*1^ BSA, and 5% glycerol at 30 °C for 2 h. After amplification, samples were washed twice with DEPC-PBSTR (20 min each).

For fluorescence detection, samples were washed three times with 10% formamide in 2× SSC (5 min each) and counterstained with DAPI. Imaging was performed using a Leica TCS SP8 confocal microscope with channels for DAPI, Alexa Fluor 488, Alexa Fluor 546, Alexa Fluor 594, and Alexa Fluor 647. Images were acquired using LAS X software (v3.7.2.22383; Leica).

For multi-round sequencing, fluorescence signals were removed after each imaging cycle by incubating samples twice with stripping buffer (60% formamide in 2× SSC) at room temperature. Subsequent sequencing-by-ligation cycles were performed by adding fresh sequencing mixtures.

### Point mutation detection by padlock probe-based FISH

For point mutation detection, MDCK cells were seeded in confocal culture dishes at 1.5 × 10^6^ cells per dish and cultured for 24 h. Cells were washed with PBS and infected with viruses at a multiplicity of infection (MOI) of 0.5 pfu/cell. At 6 h post-infection, cells were fixed with 4% paraformaldehyde. All experiments involving viruses were conducted in biosafety level 2 laboratories.

Following fixation, cells were washed three times with PBSTR (PBS supplemented with 0.1% Tween-20 and 0.1 U *µ*l^*−*1^ recombinant RNase inhibitor (RI) (Vazyme, R301)) for 5 min each. Cells were permeabilized with pre-cooled methanol (−20 °C) and incubated at −80 °C for 15 min, followed by treatment with 0.1 M HCl at room temperature for 5 min. Samples were then sealed in Secure-Seal hybridization chambers (Grace, 621505) for subsequent fluorescence in situ hybridization (FISH).

FISH probes were designed as padlock-like probes enabling signal amplification via rolling circle amplification (RCA). Probe P1 anchors the probe complex to the target RNA, while probes P2 and P3 hybridize across the mutation site, allowing allele-specific detection only when perfect sequence complementarity is achieved.

Prior to hybridization, probes P2 and P3 were phosphorylated at the 5^*′*^ end using T4 polynucleotide kinase (Vazyme, N102-01). For hybridization, probe P1 (400 nM) and equimolar phosphorylated probes P2 and P3 were prepared in hybridization buffer (2× SSC; Sangon, B548109) supplemented with 10% formamide (Sangon, A100606), 0.1% Tween-20 (Sigma-Aldrich, D8906), and 20 mM ribonucleoside–vanadyl complexes (Beyotime, R0108), followed by overnight incubation at 37 °C. Samples were washed twice with PBSTR for 20 min each, followed by one wash with 4× SSC in PBSTR.

For ligation, a reaction mixture containing SplintR ligase (1 U *µ*l^*−*1^; NEB, M0375L), T4 DNA ligase (1 U *µ*l^*−*1^; NEB, M0202L), 1 ×buffer, and RNase inhibitor (0.2 U *µ*l^*−*1^) was added and incubated at 37 °C for 2 h. Samples were then washed three times with PBSTR for 10 min each.

Rolling circle amplification (RCA) was performed using Phi29 DNA polymerase (1 U *µ*l^*−*1^; Vazyme, N106-01) in RCA buffer supplemented with 0.25 *µ*M dNTPs, 0.2 *µ*g *µ*l^*−*1^ BSA, and 5% glycerol at 30 °C for 6 h. Samples were subsequently washed twice with PBST for 10 min each.

For detection, samples were incubated in detection buffer (2× SSC containing 100 nM Alexa Fluor 488-labeled 627E detection probe and 100 nM Alexa Fluor 647-labeled 627K detection probe in PBS) at 37 °C for 2 h. After three washes with PBST (5 min each), images were acquired using a Nikon AX R NSPARC Confocal Microscope System equipped with a ×60 oil-immersion objective.

### DNA-FISH detection of herpesviruses

DNA-FISH was performed to detect feline herpesvirus(FHV), pseudorabies virus (PRV), and herpes simplex virus type 1 (HSV-1) in infected cells. Genome-tiling probe sets were designed based on the full viral genomes, comprising approximately 2000–3000 probes per virus (150 bp each), achieving high genome coverage. Probes were generated via amplification from probe libraries followed by in vitro transcription and reverse transcription, during which fluorescent nucleotides were incorporated to enable direct labeling.

For infection models, FHV was used to infect CRFK cells, while PRV and HSV-1 were used to infect Vero cells. Cells were cultured until approximately 70–80% cytopathic effect was observed, then collected and washed with PBS. Cells were permeabilized using pre-warmed hypotonic KCl solution at 37 °C for 30 min and subsequently fixed with methanol/acetic acid (3:1) at room temperature for 20 min.

For hybridization, fixed cells were dropped onto glass slides, dehydrated through a graded ethanol series (70%, 85%, and 100%), and air-dried. Probe hybridization mixture was applied, followed by denaturation at 82 °C for 10 min and hybridization at 50 °C for 1–2 h. After hybridization, slides were washed in pre-warmed 0.3% NP-40/0.4× SSC at 68 °C for 4 min.

Nuclei were counterstained with DAPI (1:1000 dilution) for 4 min, followed by mounting with antifade medium. Fluorescence signals were visualized using a fluorescence microscope. The presence and localization of viral genomes were determined based on virus-specific fluorescence signals, enabling qualitative and spatial detection of FHV, PRV, and HSV-1 infection.

## Supplementary Figures

**Fig. S1:**
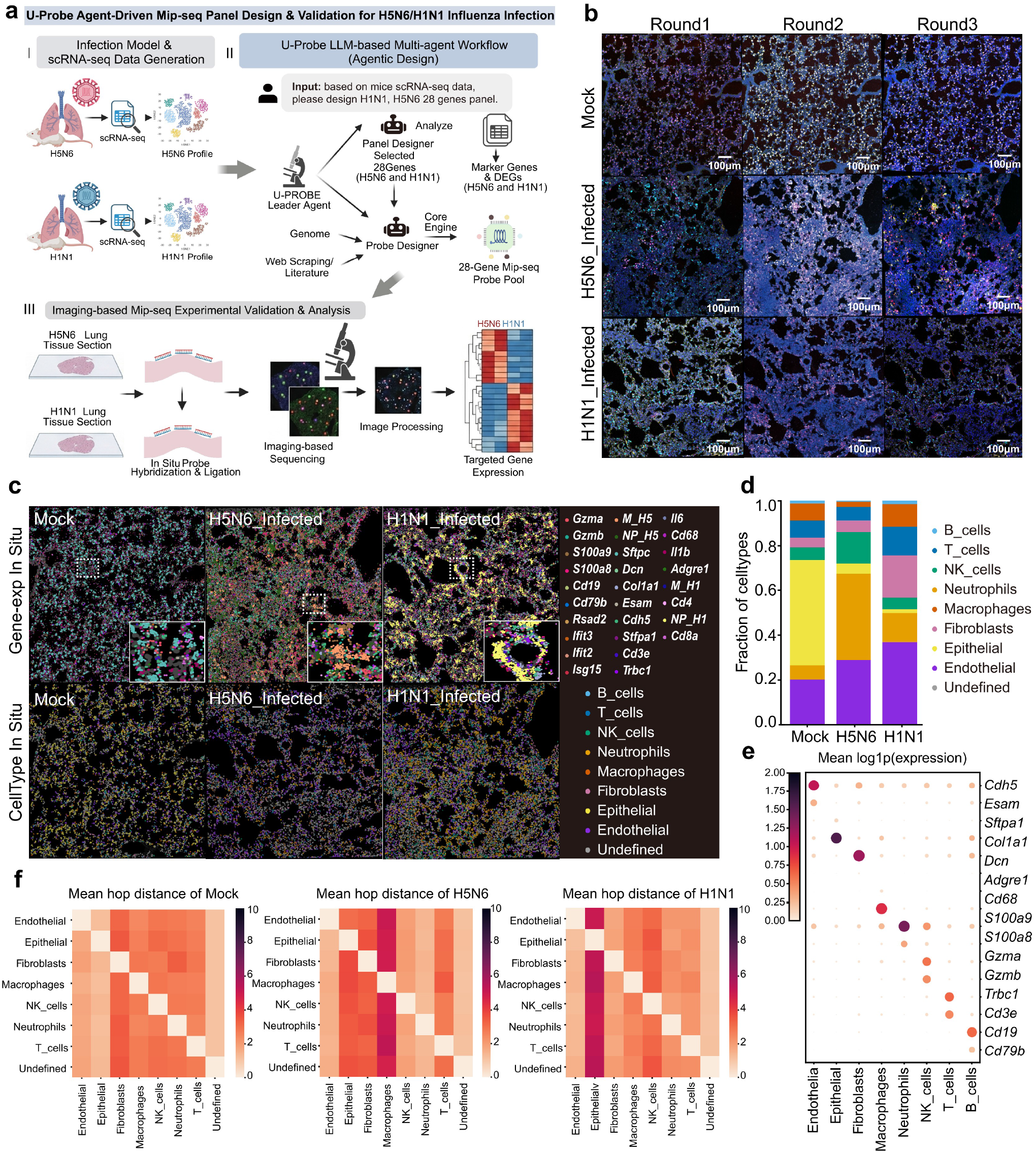
U-Probe Agent-driven MiP-seq panel design and validation for H5N6/H1N1 influenza infection. **a**, Overview of the agentic design workflow. Starting from a mouse lung scRNA-seq infection model, the U-Probe Agent executes a multi-step pipeline: analyzing input data, selecting marker genes, and generating an optimized MiP-seq probe panel through conversational interaction. Designed probes are then applied for imaging-based spatial transcriptomics validation. **b**, Representative MiP-seq spatial fluorescence imaging of mock, H5N6-infected, and H1N1-infected mouse lung tissue sections (Supplementary Table S3). **c**, Spatial gene expression maps and cell type distributions across conditions, showing the spatial localization of distinct cell populations. **d**, Cell type composition analysis across conditions, revealing differential immune cell infiltration upon viral infection. **e**, Dot plot of marker gene expression across cell types, where dot size represents the proportion of expressing cells and color intensity indicates expression level. **f**, Heatmap of average hop distance^5^ between cell types and all other cell types under mock, H5N6-infected, and H1N1-infected conditions.

**Fig. S2:**
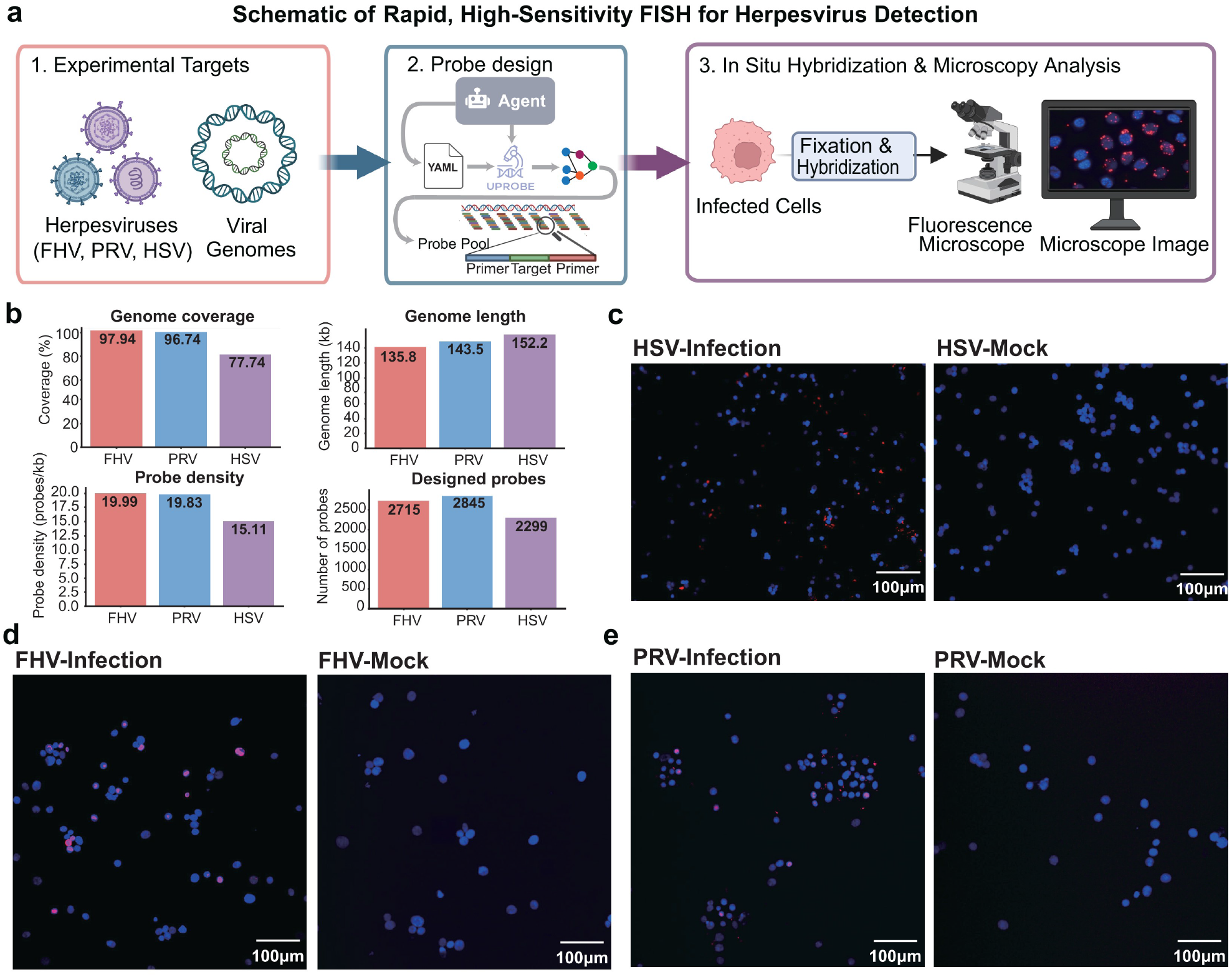
Rapid, high-sensitivity DNA-FISH for herpesvirus detection using U-Probe-designed probes. **a**, Schematic of the experimental workflow. Three herpesviruses—feline herpesvirus (FHV), pseudorabies virus (PRV), and herpes simplex virus (HSV)—were selected as targets. U-Probe Agent generated YAML configurations and designed genome-tiling probe pools, which were applied to infected cells for in situ hybridization and fluorescence microscopy analysis. **b**, Summary statistics of the designed probe sets, including genome length, genome coverage, number of probes, and probe density (probes/kb) for each virus. **c–e**, Representative fluorescence microscopy images of HSV-infected versus mock cells (**c**), FHV-infected versus mock cells (**d**), and PRV-infected versus mock cells (**e**). Viral FISH signals are shown in red/magenta; nuclei are stained in blue (DAPI).

**Fig. S3:**
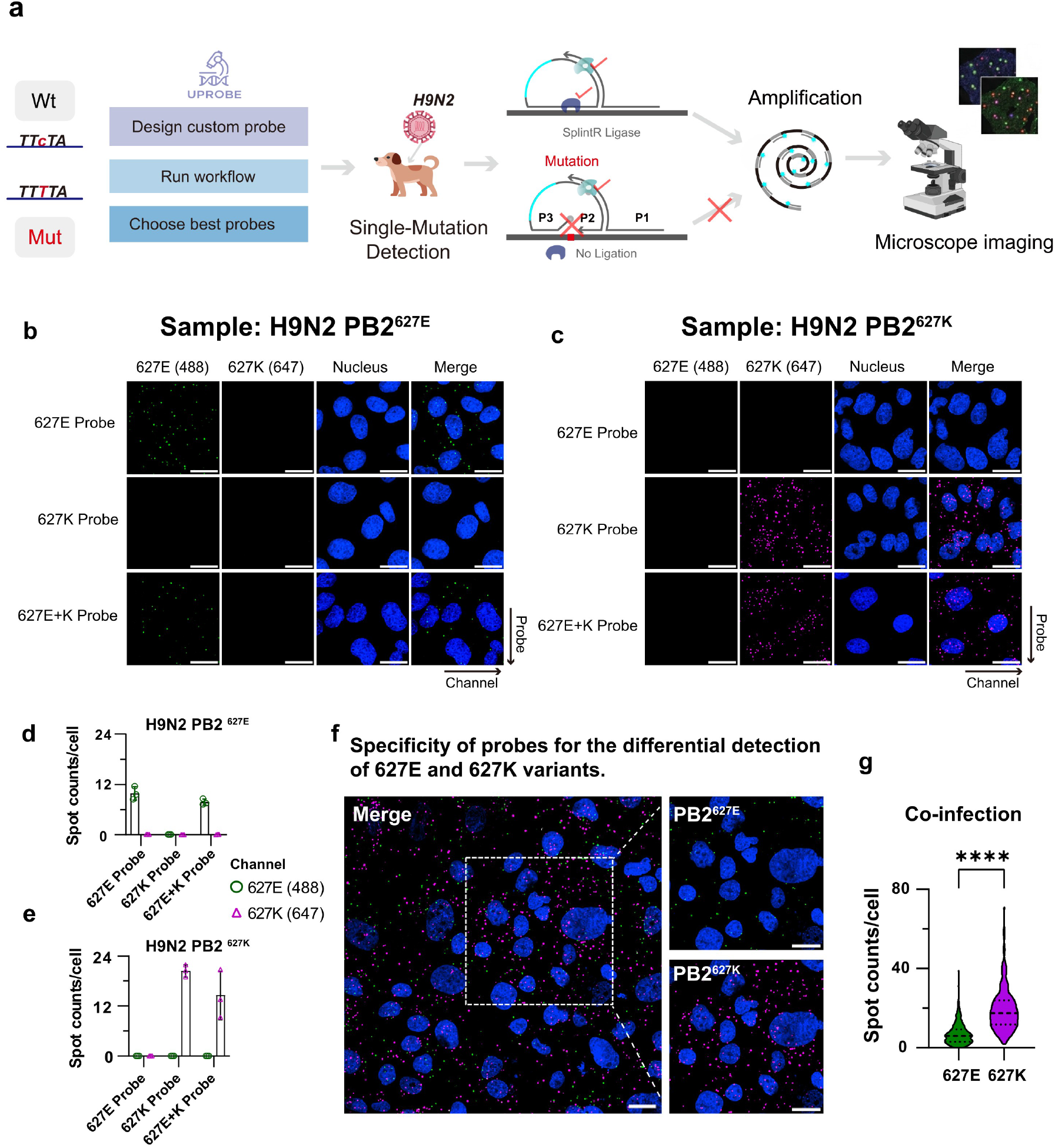
Novel RCA-based probe design for single-nucleotide mutation detection in H9N2 avian influenza virus. **a**, Schematic of the U-Probe-assisted workflow. Users define custom probe structures via U-Probe for single-mutation detection of the PB2 627E/K polymorphism in H9N2 virus. Circle probes with target-complementary arms are designed to discriminate between wild-type (627E) and mutant (627K) alleles via SplintR Ligase-mediated ligation, followed by rolling circle amplification (RCA) and fluorescence microscopy imaging. **b**,**c**, Representative fluorescence images of cells infected with H9N2 PB2^627E^ (**b**) and H9N2 PB2^627K^ (**c**), hybridized with 627E-specific, 627K-specific, or combined (627E+K) probes. **d**,**e**, Quantification of FISH spot counts per cell for PB2^627E^-infected (**d**) and PB2^627K^-infected (**e**) samples, confirming allele-specific detection. **f**,**g**, Co-infection detection: cells simultaneously infected with both PB2^627E^ and PB2^627K^ strains, demonstrating multiplexed single-mutation discrimination within the same cell.

**Fig. S4:**
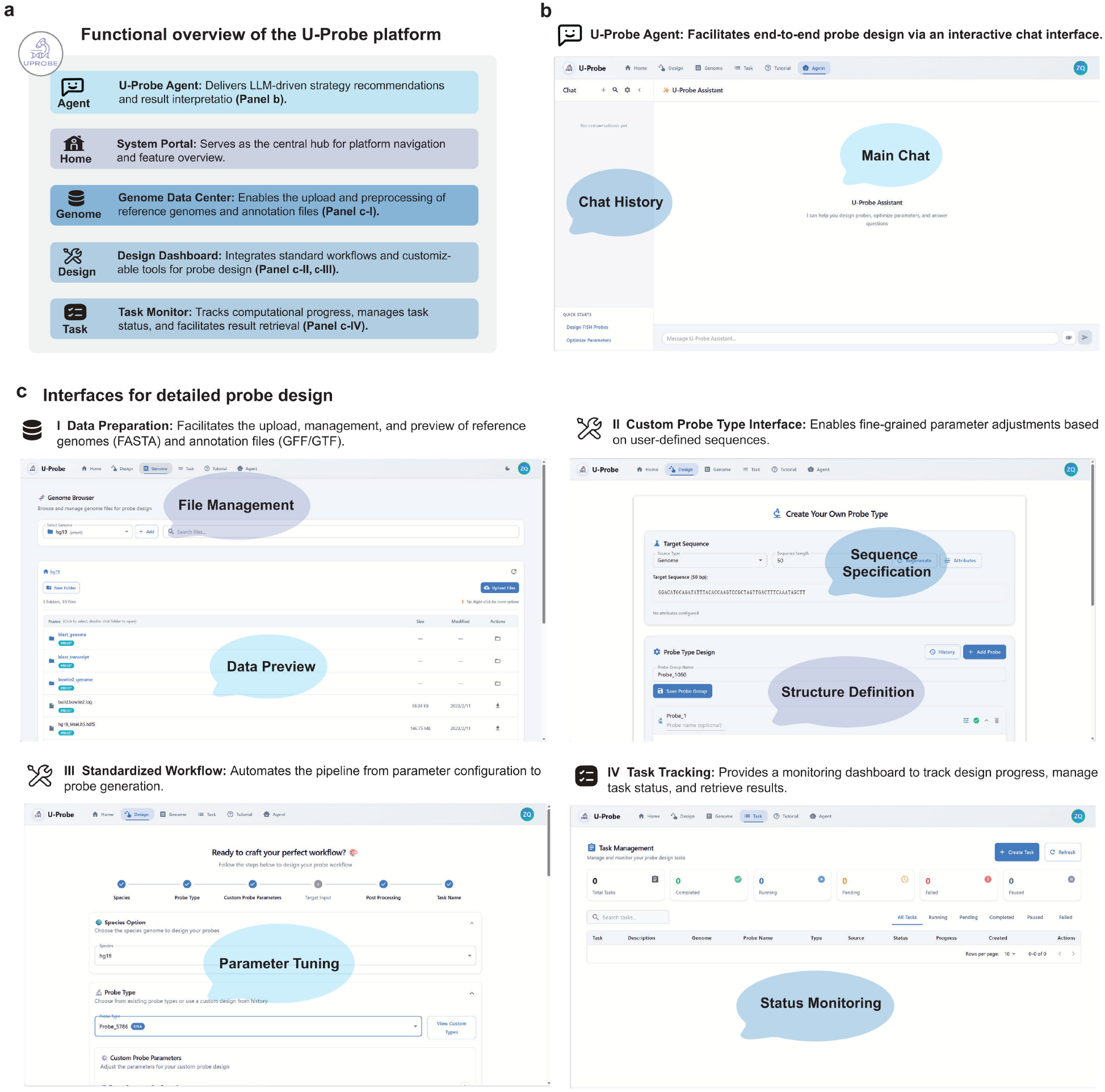
U-Probe Web platform interface. **a**, Functional overview of the U-Probe Web platform. The system comprises five modules: System Portal for navigation, Design Dashboard for standard workflows and customizable probe design tools, Genome Data Center for reference genome and annotation file management, Task Monitor for tracking computational progress, and U-Probe Agent for LLM-driven conversational design. **b**, The U-Probe Agent interface, enabling end-to-end probe design through an interactive chat with support for conversation history management. **c**, Detailed interfaces for probe design workflows: (I) Data Preparation module for uploading, managing, and previewing reference genomes (FASTA) and annotation files (GFF/GTF); (II)Custom Probe Type Interface for defining user-specified probe structures with fine-grained sequence and parameter control; (III) Standardized Workflow for automated pipeline execution from parameter configuration to probe generation; (IV) Task Tracking dashboard for monitoring design progress, managing task status, and retrieving results.

## Supplementary Tables

**Table S1:** Design constraints for advanced FISH and spatial omics modalities.

| Method | Description | Target | Probe Architecture | $T_m$<br>(°C) | GC<br>(%) | Secondary<br>Structure<br>Constraints |
| --- | --- | --- | --- | --- | --- | --- |
| **RAEFISH**<sup>6</sup> | Sequencing-free whole-transcriptome imaging via reverse padlock amplicons. | RNA, gRNA | Reverse-padlock | ~60–65<sup>a</sup> | 40–60 | No junction hairpins for ligase access. |
| **Oligo-LiveFISH**<sup>7</sup> | CRISPR-based genomic dynamics imaging in living cells. | DNA (Live) | crRNA/sgRNA | ~37<sup>b</sup> | 40–60 | Maintain dCas9-scaffold; avoid spacer hairpins. |
| **PRISM**<sup>8</sup> | Multiplexed RNA FISH via color-intensity profiling. | RNA | Color-barcoded | ~70 | 45–55 | Filter crosstalk and complex folding. |
| **TDDN-FISH**<sup>9</sup> | Signal-enhanced spatial detection via DNA nanostructure assembly. | DNA/RNA | Nanostructures | ~65–70 | 40–60 | Restrict unintended folding for self-assembly. |
| **OpenFISH**<sup>10</sup> | Open-source framework for high-throughput probe design. | DNA/RNA | Primary + Readout | ~65–75 | 40–60 | Avoid self-complementarity for specificity. |
| **MiP-seq**<sup>11</sup> | Subcellular multi-omics via pairwise sequencing of MIPs. | DNA/RNA | MIPs (Pairwise) | ~60–65 | 40–60 | Linkers must not interfere with targeting arms. |
| **ExSeq**<sup>12</sup> | Expansion sequencing; physical magnification with *in situ* sequencing. | RNA | Padlock / Hexamers | ~60 | ~50 | No steric hindrance for circularization. |
| **seqFISH+**<sup>13</sup> | Sequential FISH with pseudocolor coding for multiplexing. | DNA/RNA | U-shaped probe | ~65–70 | 45–65 | Independent binding of encoding probes. |
| **STARmap**<sup>14</sup> | Hydrogel-tissue chemistry for 3D transcriptomic readout. | RNA | SNAIL probes | 40–60<sup>c</sup> | 45–60 | Stable SNAIL loop for RCA initiation. |
| **MERFISH**<sup>15</sup> | Error-robust FISH using combinatorial barcoding and correction. | RNA | Encoding + Readout | ~65–75 | 45–65 | No strong structures ($\Delta G < -5$ kcal/mol). |

**Note:** ~, approximate value; –, recommended range. All $T_m$ assume standard buffer (e.g., 2× SSC). <sup>a</sup>Ligation junction $T_m$ .
<sup>b</sup>Physiological temperature for cell viability. <sup>c</sup>Enzymatic reaction temperature. **Abbreviations:** MIP, molecular inversion probe; RCA, rolling circle amplification; $T_m$ , melting temperature.

**Table S2:** Benchmarking U-Probe against existing FISH probe design tools. Tools are listed in reverse chronological order (2025–2002). U-Probe integrates diverse alignment engines with highly modular probe architectures, facilitating the development of emerging spatial omics and FISH-based imaging technologies.

| Tool | Year | Target Type | Interface | Alignment Engine | Property Filtering | Custom Arch.<sup>a</sup> | *De novo* Supp.<sup>b</sup> |
| --- | --- | --- | --- | --- | --- | --- | --- |
| **U-Probe** | 2025 | DNA/RNA | Agent, Web, CLI | BLAST, Bowtie2, MMseqs2, k-mer | $T_m$, GC, Fold score, Self-match, Mapped genes, K-mer counts, Mapped sites | **Yes** | **Yes** |
| Tigerfish<sup>16</sup> | 2024 | DNA | CLI | Bowtie2 | $T_m$, GC, k-mer counts | Yes | Yes |
| PaintSHOP<sup>17</sup> | 2021 | DNA/RNA | Web | ML / k-mer | Database-specific | No | No<sup>c</sup> |
| Chorus2<sup>18</sup> | 2021 | DNA | CLI | BWA, k-mer | Parameterized ($T_m$, k-mer) | No | Yes |
| ProbeDealer<sup>19</sup> | 2020 | DNA/RNA | GUI, CLI | Bowtie | Basic parameters | Yes | Yes |
| iFISH<sup>20</sup> | 2019 | DNA | Web, CLI | Bowtie, pre-computed | Parameterized | Yes | Yes<sup>d</sup> |
| OligoMiner<sup>21</sup> | 2018 | DNA/RNA | CLI | Bowtie2, BLAST | Parameterized | Limited<sup>e</sup> | Yes |
| mathFISH<sup>22</sup> | 2011 | RNA | Web | Thermodynamics models | Advanced models | No | N/A |
| PROBER<sup>23</sup> | 2006 | DNA | GUI, CLI | Suffix array / MerMatch | $T_m$, Repeat masking | No | Yes |
| OligoArray<sup>24</sup> | 2002 | DNA/RNA | Standalone | BLAST | $T_m$, GC, Mfold | No | Yes |

<sup>a</sup>Capability to append custom readouts, barcodes, or secondary structures for spatial transcriptomics.
<sup>b</sup>Probe design from scratch using user-provided, unindexed FASTA files.
<sup>c</sup>Relies on pre-mined genomic databases.
<sup>d</sup>Requires command-line package (ifpd) for *de novo* design.
<sup>e</sup>Requires manual scripting.
**Abbreviations:** BWA, Burrows-Wheeler Aligner; CLI, command-line interface; GUI, graphical user interface; $T_m$ , melting temperature.

**Table S3:** MiP-seq barcode encoding scheme for the influenza infection panel. Each gene is assigned a unique 6-nt barcode composed of two informative positions and two degenerate (N) positions. Fluorophore wavelengths: A = 488 nm, C = 594 nm, G = 546 nm, T = 647 nm.

| Gene | Barcode | Cell Type / Function | Probe Pairs |
| --- | --- | --- | --- |
| *Cd19* | GGAANN | B cells | 3 |
| *Cd79b* | GGGGNN | B cells | 3 |
| *Esam* | CCGGNN | Endothelial | 3 |
| *Cdh5* | CCCCNN | Endothelial | 3 |
| *Sftpc* | GGCCNN | Epithelial | 3 |
| *Stfpa1* | CCTTNN | Epithelial | 3 |
| *Dcn* | GGTTNN | Fibroblasts | 3 |
| *Col1a1* | CCAANN | Fibroblasts | 3 |
| *M_H1* | NNGGGG | H1N1 viral gene | 3 |
| *NP_H1* | NNGGAA | H1N1 viral gene | 3 |
| *M_H5* | NNGGTT | H5N6 viral gene | 3 |
| *NP_H5* | NNGGCC | H5N6 viral gene | 3 |
| *Rsad2* | NNTTTT | Interferon response | 5 |
| *Ifit3* | NNTTCC | Interferon response | 5 |
| *Ifit2* | NNTTGG | Interferon response | 5 |
| *Isg15* | NNCCTT | Interferon response | 5 |
| *Cd68* | AANNAA | Macrophages | 3 |
| *Adgre1* | AANNGG | Macrophages | 3 |
| *S100a9* | AACCNN | Neutrophils | 3 |
| *S100a8* | AATTNN | Neutrophils | 3 |
| *Gzma* | AAAANN | NK cells | 3 |
| *Gzmb* | AAGGNN | NK cells | 3 |
| *Il6* | NNCCGG | Pro-inflammatory cytokine | 5 |
| *Il1b* | NNCCAA | Pro-inflammatory cytokine | 5 |
| *Cd3e* | TTAANN | T cells | 3 |
| *Trbc1* | TTGGNN | T cells | 3 |
| *Cd4* | TTTTNN | T cells | 5 |
| *Cd8a* | TTCCNN | T cells | 5 |

**References**

